## Supporting Information for "Comparative study of two Rift Valley fever virus field strains circulating in Mauritania in 2010 and 2013 reveals the high virulence of the MRU25010-30 strain isolated from camel"

Short title: Comparison of the virulence of two RVFV field strains

Mehdi Chabert [1,2,3], Sandra Lacôte[4], Philippe Marianneau[4], Marie-Pierre Confort[1], Noémie Aurine[5], Aurélie Pédarrieu[2,3], Baba Doumbia[6], Mohamed Ould Baba Ould Guéya[6], Habiboullah Habiboullah[6], Ahmed Bezeid Ould El Mamy[7], Modou Moustapha Lo[8], Jenna Nichols[9#], Vattipally B Sreenu[9], Ana da Silva Filipe[9], Marie-Anne Colle[10], Bertrand Pain[5], Catherine Cêtre-Sossah[2,3,¶], Frédérick Arnaud[1,¶] and Maxime Ratinier[1,*,¶]

[1] IVPC UMR754, INRAE, Universite Claude Bernard Lyon 1, EPHE, PSL Research University, F-69007 Lyon, France.

[2] CIRAD, UMR ASTRE, F-34398 Montpellier Cedex, France

[3] ASTRE, Univ Montpellier, CIRAD, INRAE, Montpellier, France

[4] ANSES, Virology Unit, F-69007 Lyon, France

[5] Université Lyon 1, INSERM, INRAE, Stem Cell and Brain Research Institute, U1208, USC1361, F-69675 Bron, France

[6] Direction des Services Vétérinaires, Ministère de l'élevage, Nouakchott, Mauritania

[7] ONARDEP, Nouakchott, Mauritania

[8] ISRA-LNERV, Dakar, BP 2057 Dakar Hann, Senegal

[9] MRC-University of Glasgow Centre for Virus Research, Glasgow, United Kingdom.

[10] ONIRIS-UMR 703-PAnTher INRAE/Oniris, F-44307 NANTES Cedex 3, France

### Current address: School of Biodiversity, One Health and Veterinary Medicine, University of Glasgow, Glasgow, United Kingdom

¶ These authors contributed equally to this work

### Supporting information

**Table S1: Mapping statistics of the high throughput sequencing data.**

| | | L segment | | M segment | | S segment | |
|---|---|---|---|---|---|---|---|
| Strain | Total reads | Mapped reads | Average depth | Mapped reads | Average depth | Mapped reads | Average depth |
| MRU25010-30 | 2067461 | 224815 (10.87%) | 9332 | 42895 (2.07%) | 3062 | 53173 (2.57%) | 8509 |
| MRU2687-3 | 1598621 | 125040 (7.82%) | 5160 | 151253 (9.46%) | 10448 | 63700 (3.98%) | 9926 |
| ZH548 | 905582 | 35163 (3.88%) | 1353 | 39421 (4.35%) | 2614 | 8063 (0.89%) | 1199 |

**Table S2: Amino acid substitutions observed between the consensus sequences of MRU25010-30, MRU2687-3 and ZH548 strains.**

| Segment | Protein | Position | MRU25010-30 | MRU2687-3 | ZH548 |
|---|---|---|---|---|---|
| S | NSs | 23 | I | I | F |
| | | 90 | V | I | I |
| | | 111 | V | I | I |
| | | 167 | V | V | A |
| | | 217 | A | A | V |
| | | 242 | V | V | I |
| | | 245 | V | I | I |
| | | 262 | V | A | V |
| | N | 159 | E | E | G |
| M | NSm | 42 | R | G | G |
| | | 126 | V | I | I |
| | Gn | 232 | Q | Q | L |
| | | 269 | V | I | V |
| | | 360 | D | N | D |
| | | 384 | K | T | T |
| | | 492 | I | V | V |
| | | 566 | G | G | D |
| | | 595 | V | I | I |
| | | 605 | K | R | R |
| | | 615 | K | R | R |
| | | 631 | V | V | I |
| | | 659 | A | V | V |
| | Gc | 739 | D | E | E |
| | | 747 | I | I | L |
| | | 1059 | T | T | S |
| L | L | 23 | Y | Y | F |
| | | 120 | M | T | M |
| | | 157 | D | G | G |
| | | 177 | E | D | E |

| | | | | | |
|---|---|---|---|---|---|
| | | 249 | K | R | R |
| | | 278 | N | N | S |
| | | 288 | V | A | A |
| | | 302 | I | V | V |
| | | 336 | I | V | V |
| | | 350 | K | R | R |
| | | 406 | V | M | M |
| | | 407 | D | G | G |
| | | 411 | G | S | S |
| | | 470 | N | N | S |
| | | 493 | R | K | R |
| | | 663 | T | T | A |
| | | 840 | I | V | I |
| | | 1333 | I | V | V |
| | | 1698 | I | V | V |
| | | 1751 | M | I | I |
| | | 1760 | V | I | V |
| | | 1852 | H | Y | H |
| | | 1926 | R | K | R |
| | | 1960 | E | D | D |
| | | 1984 | N | N | D |
| | | 2033 | T | A | A |

Amino acid substitutions are classified by viral segment and related protein. Their position was determined relative to the known start codon. Note that the numbering of M segment proteins (NSm, Gn and Gc) starts from AUG1 used to translate p78. Amino acid residues can be classified in four groups based on their polarity (non-polar, polar with no charge on R group, polar with negative charge on R group and polar with positive charge on R group). Amino acid residues conserved between MRU25010-30 and MRU2687-3 are uncoloured. Non-conserved amino acid residues between these two strains but identical between MRU25010-30 and ZH548 strains are coloured in light grey. Non-conserved amino acid residues between MRU25010-30 and MRU2687-3 and from the same group are coloured in grey. Non-conserved amino acid residues and from a different group are coloured in dark grey.

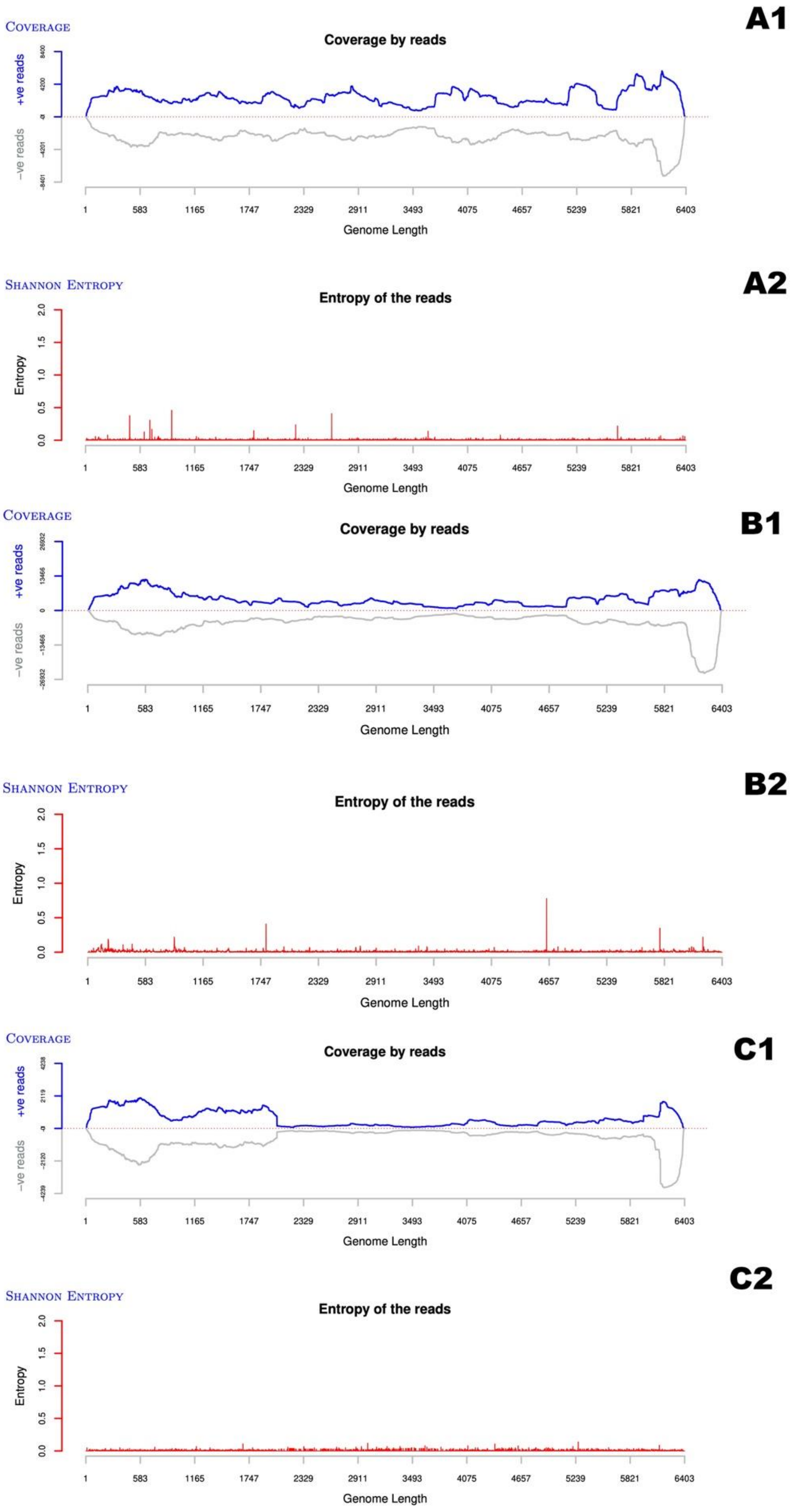


**Figure S1: Coverage and Shannon entropy plots for L segments.** Coverage (1) and Shannon entropy (2) for L segments of MRU2687-3 (A), MRU25010-30 (B) and ZH548 (C).

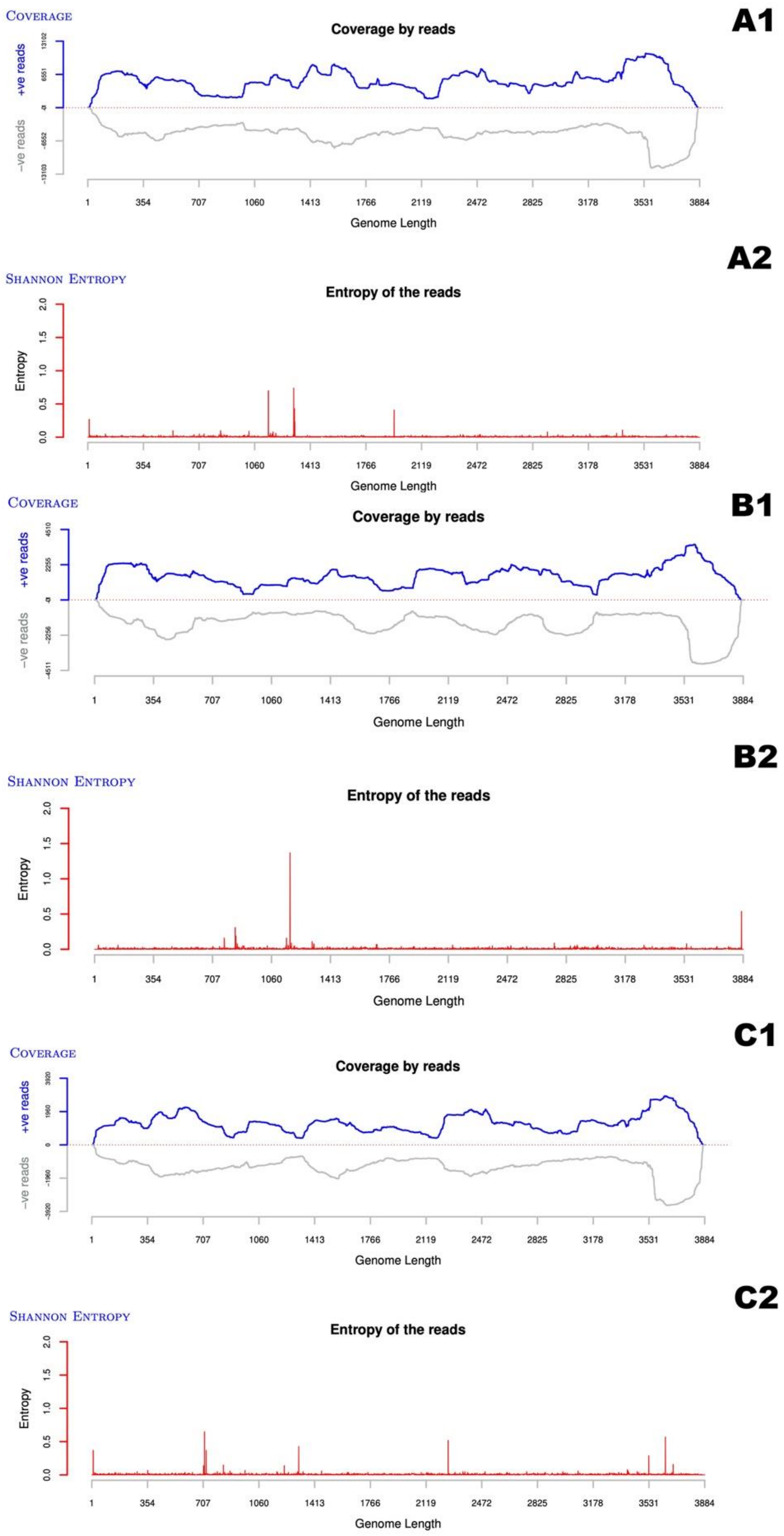


**Figure S2: Coverage and Shannon entropy plots for M segments.** Coverage (1) and Shannon entropy (2) for M segments of MRU2687-3 (A), MRU25010-30 (B) and ZH548 (C).

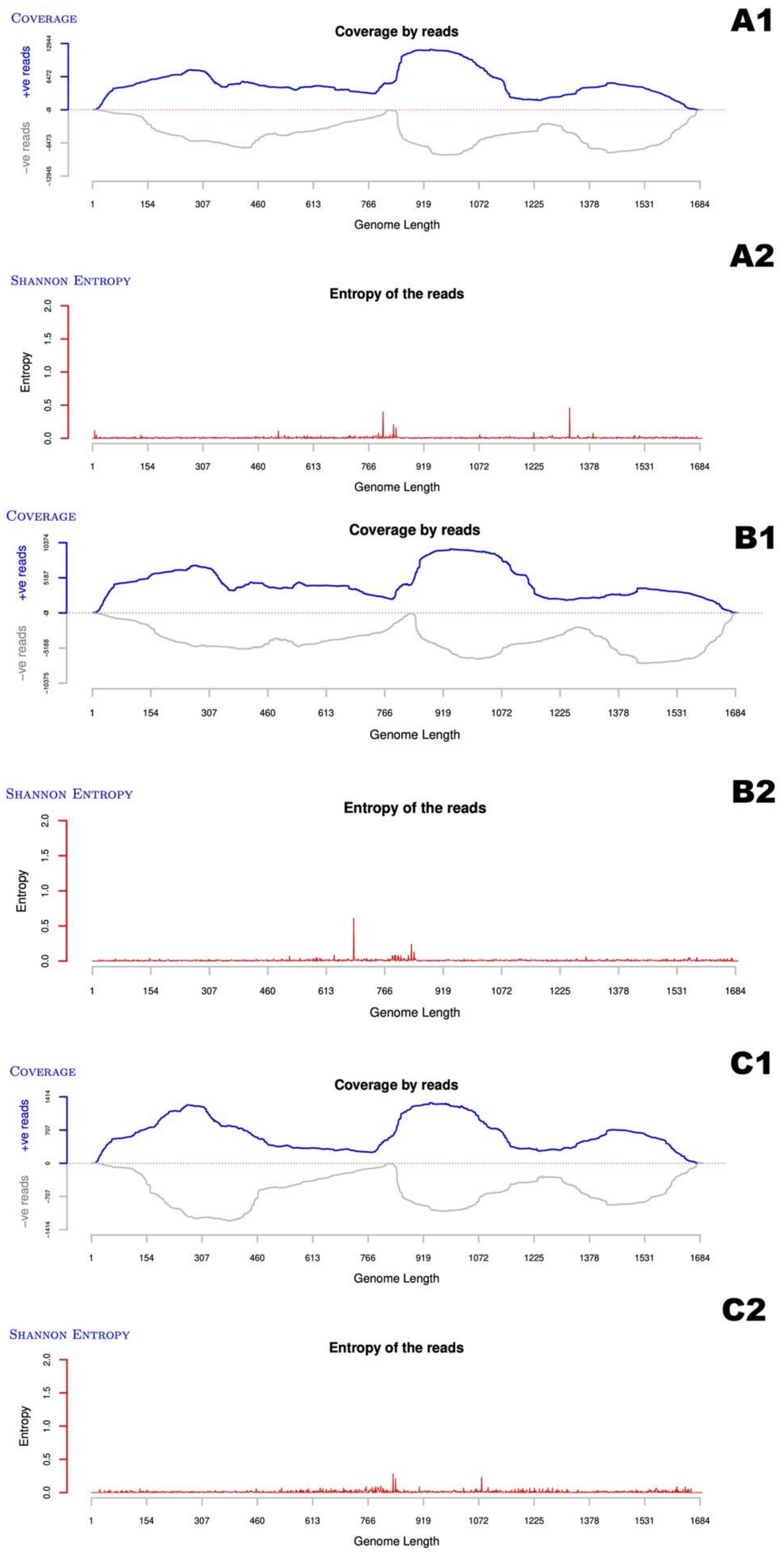


**Figure S3: Coverage and Shannon entropy plots for S segments.** Coverage (1) and Shannon entropy (2) for S segments of MRU2687-3 (A), MRU25010-30 (B) and ZH548 (C).

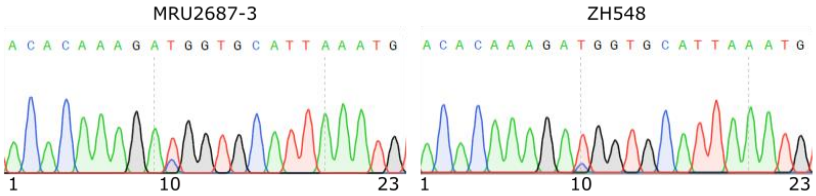


**Figure S4 : DNA sequencing chromatograms of RACE-PCR products amplified from 5'UTR of anti-genomic M segments.** Nucleotides at position 10 is either an uracile (U) or a cytosine (C) in MRU2687-3 and ZH548 strains.

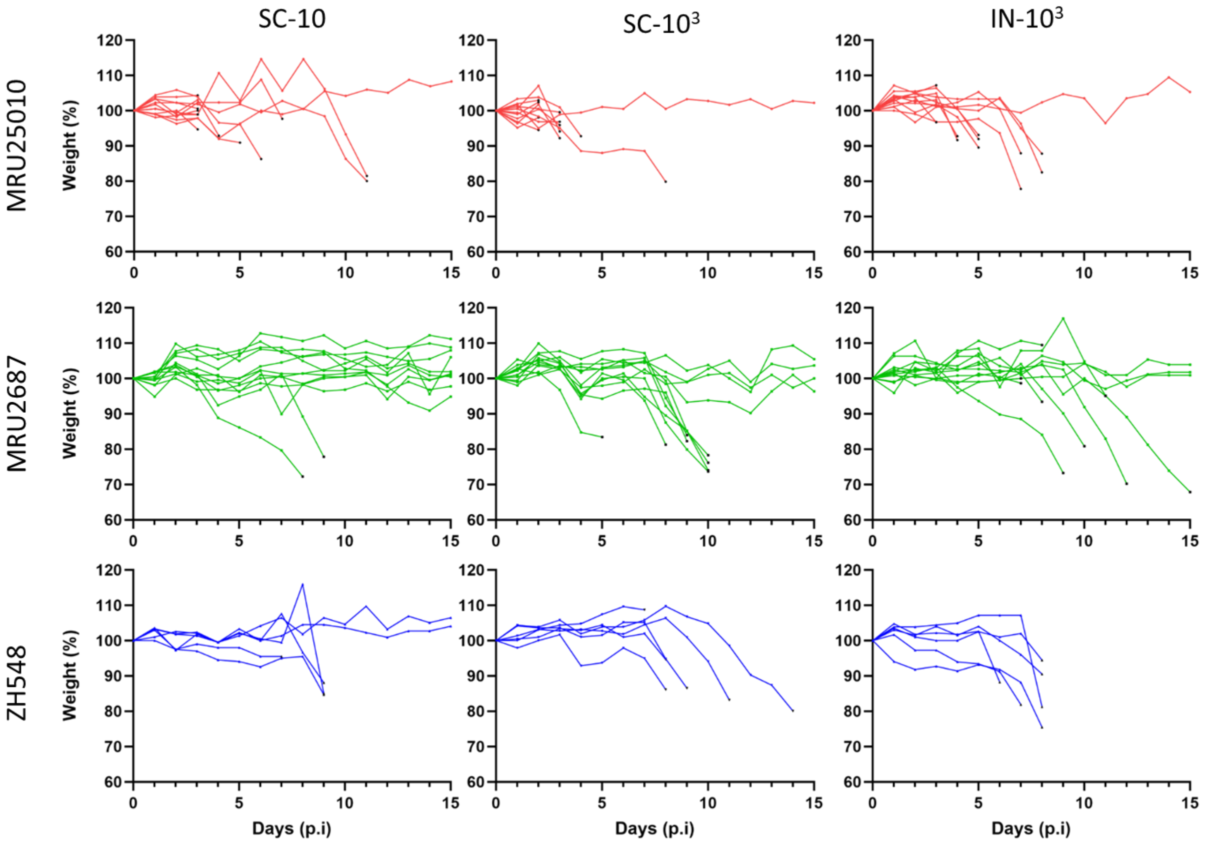


**Figure S5 : Effect of RVFV infection on BALB/c mice body weight.** Mice have been weighted before infection and then every day during the course of the experiment. Weight percentage was calculated relative to that at Day 0. Black dots represent the last measure recorded before the death of the animal.

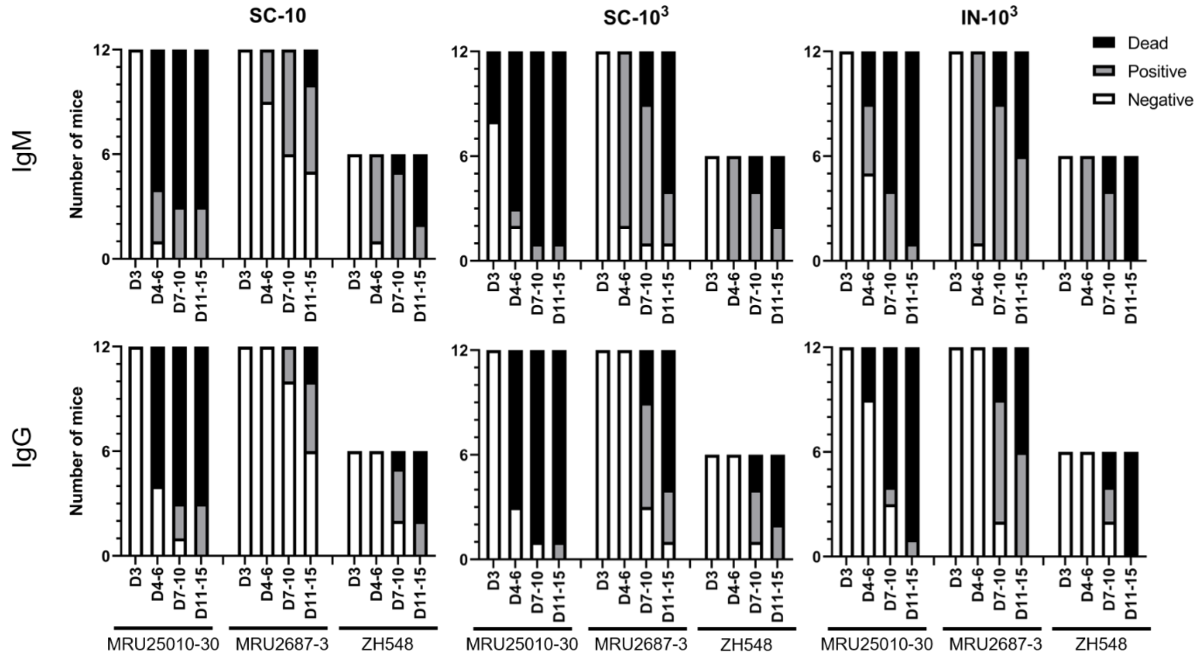


**Figure S6: Kinetics of seroconversion of mice infected with MRU25010-30, MRU2687-3 and ZH548 strains.** Anti-RVFV antibodies within mice sera were detected using in-house IgM and IgG ELISAs (Enzyme-Linked ImmunoSorbent Assay), as previously described [41,48]. Briefly, RVFV antigens were prepared from RVFV MP12 strain infected VeroE6 cells (MOI=0,01; 2 days post-infection). For IgM detection, 96 -well plates (Nunc MaxisorpTM, Thermo Fisher Scientific) were coated with rabbit anti-mouse IgM antibody (100 µL/well, 1:400 dilution; Sigma, SAB3701197) and incubated with 100 µL/well of 1:100 dilution of mice sera. RVFV antigens were subsequently detected with hyperimmunised sera from hamster infected with ZH501 strain and Goat anti-Hamster IgG (H+L)-HRP (Horseradish Peroxidase) conjugated antibody. For IgG detection, plates were coated with RVFV antigens, further incubated with 100 µL/well of 1:100 dilution of mice sera and subsequently with HRP-conjugated rabbit anti-mouse IgG (whole molecule, 1:5000, Sigma, A9044). HRP enzymatic activity was revealed using TMB substrate (Thermo Fischer Scientific). Optical density at 450 nm (OD450) was measured using a TECAN microplate reader. ELISA measurement of IgM and IgG antibodies of mice infected with field strains or ZH548. The sera were collected at days 3, 4-6, 7-10 and 11-15 pi. At the indicated day, black bars represent dead mice, grey bars seroconverted mice and white bars mice with non-detectable IgM or IgG antibodies.
